# Synchronous theta networks characterize successful memory retrieval

**DOI:** 10.1101/2024.07.12.603286

**Authors:** Aditya M. Rao, Riley D. DeHaan, Michael J. Kahana

## Abstract

Memory retrieval activates regions across the brain, including not only the hippocampus and medial temporal lobe (MTL), but also frontal, parietal, and lateral temporal cortical regions. What remains unclear, however, is how these regions communicate to organize retrieval-specific processing. Here, we elucidate the role of theta (3–8 Hz) synchronization, broadly implicated in memory function, during the spontaneous retrieval of episodic memories. Analyzing a dataset of 382 neurosurgical patients (213 male, 168 female, 1 unknown) implanted with intracranial electrodes who completed a free recall task, we find that synchronous networks of theta phase synchrony span the brain in the moments before spontaneous recall, in comparison to periods of deliberation and incorrect recalls. Hubs of the retrieval network, which systematically synchronize with other regions, appear throughout the prefrontal cortex and lateral and medial temporal lobes, as well as other areas. Theta synchrony increases appear more prominently for slow (3 Hz) theta than for fast (8 Hz) theta in the recall–deliberation contrast, but not in the encoding or recall–intrusion contrast, and theta power and synchrony positively correlate throughout the theta band. These results implicate diffuse brain-wide synchronization of theta rhythms, especially slow theta, in episodic memory retrieval.

**Significance Statement:** Analyzing intracranial recordings from 382 subjects who completed an episodic free recall experiment, we study the brain-wide theta synchrony effects of memory retrieval. The literature has not previously described the whole-brain regional distribution of these effects nor studied them with respect to intrusions. We show that a whole-brain theta synchrony effect marks the recall accuracy contrast, that distributed synchronous hubs constitute a whole-brain retrieval network, and that theta synchrony in the successful encoding, successful retrieval, and recall accuracy contrasts correlates positively with theta power increases at a region. These findings advance our understanding of the role and localization of theta synchrony effects during human memory retrieval.

## Introduction

Complex cognitive processes such as the formation and retrieval of declarative memories de-pend on the coordination of regions across the brain for information processing. Consequently, characterizing the patterns of connectivity that distinguish successful cognition from unsuccessful cognitive effort is vital to understanding the neural computations behind human memory.

The electrical activity of neuronal assemblies generates oscillatory rhythms that can become phase-locked when coupled (Lachaux et al., 1999). Phase-locking thus lends itself to a measure of functional connectivity, and intracranial recordings reveal these transient connections with high temporal and spatial resolution (Fell and Axmacher, 2011). Phase-locking of slow oscillations, called the theta rhythm (3–8 Hz), has proven a robust correlate of successful memory in various experimental paradigms, including recognition memory, spatial memory, and free recall, and memory functions, such as encoding and retrieval (Buzsáki, 2002; Siapas et al., 2005; Benchenane et al., 2010; Rutishauser et al., 2010; Sigurdsson et al., 2010; Addante et al., 2011; Solomon et al., 2019; Roux et al., 2022). Manipulations of theta phase synchrony have been linked to the formation of mnemonic associations (Clouter et al., 2017), and post-encoding theta entrainment from audiovisual stimuli has been found to enhance human memory performance (Roberts et al., 2018). Recent studies have reported that whole-brain networks of theta phase synchronization underlie episodic memory encoding, with regional hubs that reliably capture this effect distributed throughout the brain (Solomon et al., 2017; Solomon et al., 2019).

Although prior research has thoroughly characterized macroscale theta synchrony in episodic memory encoding (Mölle et al., 2002; Fell et al., 2003; Fell et al., 2008; Clouter et al., 2017; Solomon et al., 2017; Solomon et al., 2019), its role in memory retrieval is poorly understood. Studies have implicated widespread low-frequency phase-locking in memory retrieval (Solomon et al., 2017; Solomon et al., 2019; Watrous et al., 2013), and identified a few relevant interregional connections. These studies often focus on the medial temporal lobe and hippocampus, as several decades of lesion, electrophysiology, and imaging studies have corroborated their activation in episodic memory and spatial navigation (Squire and Zola-Morgan, 1991; Eichenbaum, 2000; Wixted and Squire, 2011; Solomon et al., 2019). For example, transient phase-locking between the left medial temporal lobe and left retrosplenial cortex is linked to autobiographical memory retrieval (Foster et al., 2013). Nevertheless, previous studies have not systematically detailed how regions throughout the whole brain participate in the synchronous theta networks of memory retrieval, nor how these networks change across the 3–8 Hz band in an episodic memory paradigm, despite evidence that low and high theta frequencies represent distinct functional components of the theta band (Lega et al., 2012; Watrous et al., 2013; Goyal et al., 2020). Other properties of these networks, such as their relationship to memory-related theta power changes or to the theta connectivity that underlies the same information’s encoding, remain largely unexplored.

Here, we aim to answer four questions about the synchronous theta activity supporting successful and accurate memory retrieval in an episodic free recall paradigm. First, what are the key regional hubs and connections of the retrieval and recall accuracy networks, and what role do the MTL and hippocampus play in these networks? Second, how do theta synchrony correlates of successful memory differ across the frequencies of the theta band? Third, how do participants’ retrieval and recall accuracy networks compare with their encoding networks? Fourth, what is the relation between spectral power and phase synchrony effects of successful memory?

Analyzing hundreds of subjects’ intracranial electrophysiological recordings, we detail the distributed patterns of whole-brain theta synchrony associated with successful retrieval, both when contrasted with baseline periods of silence and with incorrect recalls. We also discover that retrieval–deliberation synchrony effects are stronger in slow theta than in fast theta and that increases in theta power at a region correlate with increases in the region’s brain-wide theta synchrony. These findings advance our understanding of the functional connectivity and electrophysiological correlates of memory retrieval.

### Experimental Procedures

#### Participants

382 patients (213 male, 168 female, 1 unknown) with pharmacoresistant epilepsy completed a delayed free recall memory task while implanted with intracranial electrodes for seizure monitoring. A number of hospitals contributed data to the project as part of a multi-site collaboration: the Hospital of the University of Pennsylvania, Thomas Jefferson University Hospitals, Freiburg University Hospital, University of Texas Southwestern Medical Center, University of Texas Health Science Center at San Antonio, the National Institute of Neurological Disorders and Stroke, Dartmouth-Hitchcock Medical Center, the Mayo Clinic, UCHealth University of Colorado Hospital, and Emory University Hospital. The electrode configurations — depth, strip, or grid — and electrode contact locations were selected according to clinical considerations alone. All patients provided informed consent to the study, and each hospital’s institutional review board approved the research protocol before study activities began.

#### Free recall memory task

Patients performed a delayed free recall memory task on a laptop computer, while EEG data were recorded from their implanted electrodes at a sampling rate between 200 and 2000 Hz, depending on the settings of the clinical system in place. The patient’s behavior in the memory task was synchronized with the EEG recording either with electrical synchronization pulses or with timing messages transmitted online between the computer that recorded the EEG signal and the computer on which the participant performed the memory task.

A full session of the memory task consisted of 12 or 25 trials identical in structure. Each trial consisted of three phases: encoding, a distractor task, and retrieval (Figure 1A). During the encoding phase, words randomly drawn from one of two wordpools of common English nouns appeared one at a time for 1600 ms on the screen, with a 750–1000 ms randomly jittered interstimulus interval, or in a variant of the experiment, 800–1200 ms. Each trial presented 12 words, or in the variant, 15. Participants then completed an arithmetic distractor task, answering problems of the form A + B + C = ??, where A, B, and C were random integers between 1 and 9. This phase lasted for at least 20 s, or 25 s in the variant, allowing participants to finish any problem that appeared on the screen during that time. Finally, participants attempted to recall words aloud for 30 s, or 45 s in the variant, from the list of words just presented during the same trial’s encoding phase. The audio recordings of the retrieval phase were then manually annotated to evaluate the content and accuracy of the vocalized responses.

**Figure 1:**
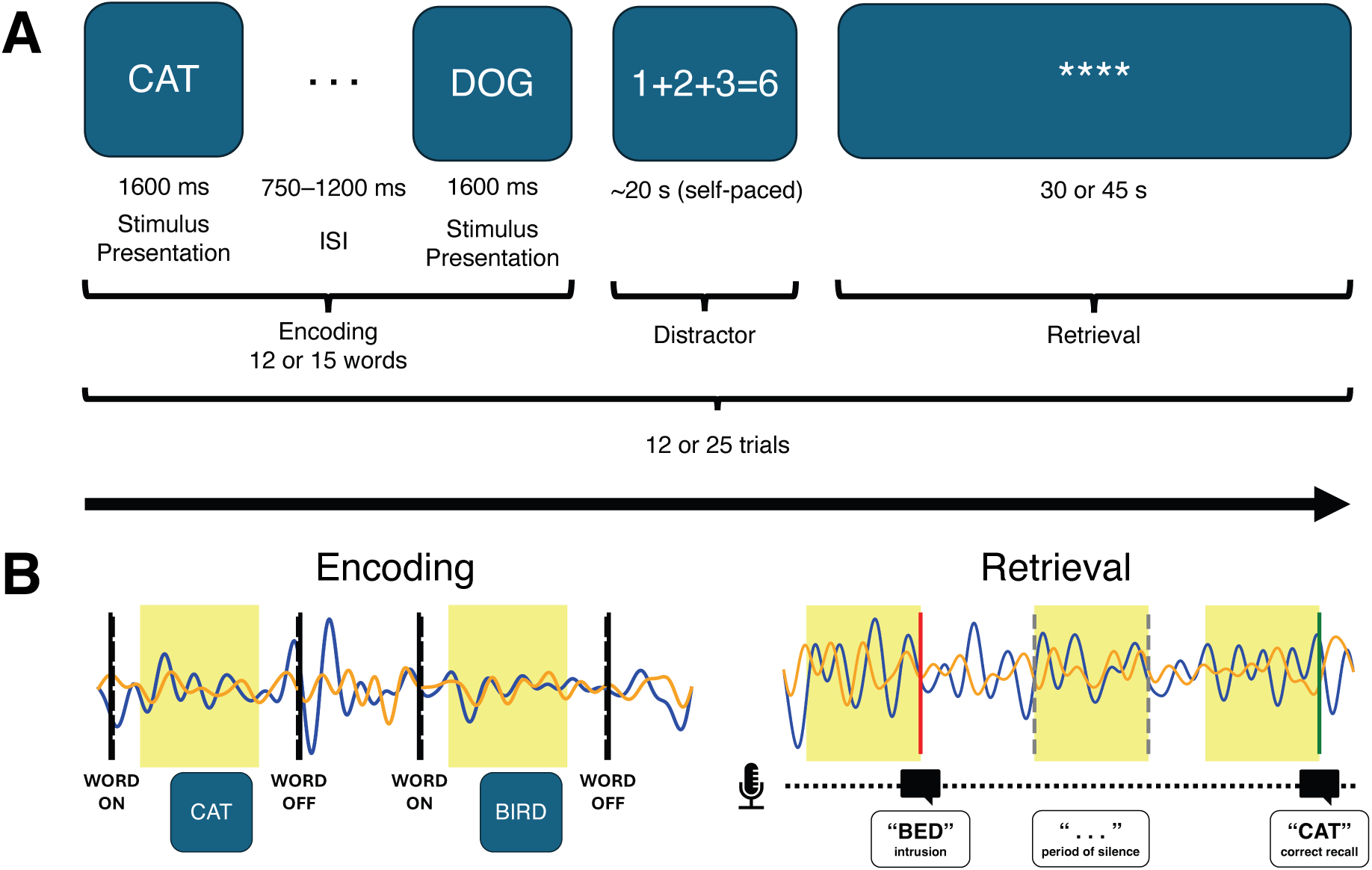
(A) Structure of free recall task. (B) Events analyzed during the encoding and retrieval phases. Analysis window highlighted in yellow.

#### Behavioral event selection

Events in the encoding and retrieval phases were classified as successful or unsuccessful. In encoding, a word presentation event was considered successful if the word was subsequently recalled during the retrieval phase of the same trial, and unsuccessful if the participant failed to recall the item. The post-presentation interval of 250–1250 ms was analyzed (Figure 1B), in order to center the 400–1100 ms window during which prior work identified the peak of the encoding-evoked theta synchrony effect (Solomon et al., 2017; Adamovich-Zeitlin et al., 2021), and to avoid spectral leakage from the preceding or following items’ presentation intervals through convolution buffers. Encoding items presented at intermediate serial positions tend to be recalled with lower probability than items presented at the beginning or end of the list. Therefore, to control for systematic differences in serial position between subsequently recalled items and not-recalled items, we matched subsequently recalled items to not-recalled items that differed in serial position by no more than 1 and analyzed only the items in these matched pairs.

In retrieval, items correctly recalled from the same trial’s encoding phase represented successful memory. Subjects recalled 31.1% ± 11.9% (mean ± SD across subjects) of presented items. We did not analyze nonverbal vocalizations. We also excluded first recalls from our analysis on the consideration that recall initiation fundamentally differs from subsequent recalls. Models of free recall (Kahana, 2012; Kahana, 2020; Howard and Kahana, 1999) support the notion that unlike the first recall, which cannot be cued by previous items, subsequent recalls in the recall phase result from associative retrieval triggered by contextual cues. Because our interest is in the cue-dependent retrieval process, and first recalls arise from a distinct process, we exclude first recalls from our analysis.

To analyze the neural activity specific to retrieval processes, we compared the pre-vocalization periods of correct recalls to periods of failed memory search. We matched the 1000–0 ms pre-vocalization analysis window of each correct recall by its vocalization onset time in the recall window to a 1000 ms period of silence in a nearby list, with preference to closer lists in view of contextual drift (Figure 1C). We matched recalls to a period of silence within 0.5 s if available or within 5 s otherwise. These periods of silence, drawn from after the first recall and before the last recall in the trial, were considered periods of unsuccessful memory search. To prevent the inclusion of vocalization artifact or neural signal associated with successful retrieval in the analyzed signal, periods of silence that began less than 1000 ms after a vocalization onset or ended less than 1000 ms before a vocalization onset were not considered. Likewise, correct recall windows that began less than 1000 ms after a vocalization onset were not analyzed to avoid vocalization artifact. We did not analyze sessions with fewer than 10 recalls that met these criteria. After applying these inclusion criteria, 34.2% ± 11.7% (mean ± SD across subjects) of subjects’ correct recalls remained in the analysis.

In the recall accuracy analysis, we analyzed correct recalls and intrusions that met the same relative timing criteria as in the retrieval analysis to avoid vocalization contamination, and matched them by vocalization onset time in the same way. Again, we did not analyze sessions with fewer than 10 pairs of correct recalls and intrusions that met these criteria. After applying these inclusion criteria, 23.4% ± 9.8% (mean ± SD across subjects) of subjects’ correct recalls remained in the analysis.

We verified that our procedures for matching events by serial position in the encoding analysis and by vocalization onset time in the retrieval and recall accuracy analyses provided similar dis-tributions of matched characteristics between successful and unsuccessful events. The difference in the average serial position between successful and unsuccessful encoding events was –0.15 ± 0.25 (mean ± SD across subjects), the difference in the average vocalization onset time in the retrieval analysis was 0.47 s ± 0.76 s (mean ± SD across subjects), and the difference in the average vocalization onset time in the recall accuracy analysis was –0.69 s ± 1.49 s (mean ± SD across subjects).

#### Anatomical localization of electrodes

After a pre-operative structural T1-weighted MRI scan was parcellated and segmented (Fischl et al., 2004), a post-operative CT scan was coregistered to it by means of Advanced Normalization Tools (ANTS) to provide coordinates for all locations in the CT scan (Avants et al., 2008). Implanted electrode contacts were manually identified and annotated according to these coordinates. An automated pipeline then mapped these coordinates to brain region labels (Dykstra et al., 2012). Two clinical neuroradiologists reviewed a subset of these labels for accuracy. For some subjects, medial temporal lobe subregions were also automatically labelled in a pre-operative T2-weighted MRI scan when available (Yushkevich et al., 2015).

80 ROIs, evenly divided between the hemispheres, constituted the nodes for network analyses. These ROIs were selected to cover the cerebrum while coinciding with standard anatomical and functional labels broadly used in the neuroimaging and functional connectivity literature, such as the Desikan-Killiany atlas (Desikan et al., 2006). All brain plots represent region locations in Montreal Neurological Institute (MNI) space.

#### EEG processing

Bipolar re-referencing was applied to all channel recordings before analysis; bipolar differencing was carried out between neighboring electrodes and neither bipolar pairs between contacts more than 20 mm apart nor channels with a flatlined signal were analyzed. All EEG recordings were resampled to 250 Hz, with 1000 ms buffers to avoid edge artifacts. For encoding events, the buffers consisted of the EEG signal adjacent to the analysis window. For recall and deliberation events, 1000 ms mirrored buffers were appended to either side of each analysis window to preclude the introduction of vocalization or motion artifact into the analyzed signal, and clipped after time-frequency decomposition. A 4th-order Butterworth bandstop filter at 58–62 Hz (United States) or 48–52 Hz (Germany) removed line noise. Morlet wavelet convolution (wave number = 5) then extracted instantaneous phase and power from the EEG recordings during the selected event windows at each integer-valued frequency in the 3–8 Hz theta band.

#### Phase synchrony analysis and functional network construction

Phase synchrony refers to the consistency across behavioral trials — periods of item encoding or memory search — of the phase lag between signals across time. To analyze phase synchrony, we computed the difference in instantaneous phase for each pair of contacts. Then, we circularly averaged the phase lag across the analysis window into five 200 ms bins, or epochs. For each pair of contacts, each frequency, and each epoch, we computed the pairwise phase consistency (PPC) (Vinck et al., 2010) for successful memory events and unsuccessful memory events separately. Finally, we averaged phase synchrony values across contacts located in the same region. The difference in pairwise phase consistency between successful and unsuccessful memory events corresponded to the network connectivity specific to improved memory function.

#### Network analyses and statistical procedures

The average ΔPPC for an ROI pair gave the connectivity value in subject-level connectivity matrices. Paired-samples *t*-tests across the subjects were used to test hypotheses about differences in theta synchrony related to successful memory in contrast with unsuccessful memory.

A region that reliably exhibits high mean synchronization to other regions in the brain is a network hub. To assess the “hubness” of a region, we averaged both successful memory and unsuccessful memory PPC values over all theta frequencies and epochs of the analysis window, and all other regions to which the test region had connections. The difference between the average successful memory PPC value and the average unsuccessful memory PPC value then yielded a within-subject “hubness score” for each region. Then, we applied a one-sample *t*-test against population mean zero to the distribution of the region’s hubness scores across those subjects for whom recordings from that region were available, and corrected the resulting *p*-values with the Benjamini-Yekutieli two-stage adaptive linear step-up procedure for FDR control at *Q*_FDR_ = 0.05 (Benjamini et al., 2006). We did not test hubness among ROIs for which less than 7 subjects’ worth of data was available. In other analyses of the MTL and hippocampus, the Holm method was applied for control of the family-wise error rate at α = 0.05 (Holm, 1979).

In our statistical significance testing in the time-frequency analysis, we used threshold-free cluster enhancement (TFCE) to account for correlations among time-frequency pixels (Smith and Nichols, 2009). To compute *p*-values, given each subject’s average PPC values for successful and unsuccessful memory events, we randomly flipped subjects’ successful and unsuccessful memory condition labels 999 times, computed the TFCE-enhanced *t*-statistics with these same 999 permutations for all time-frequency pixels, and then derived *p*-values for each time-frequency pixel’s test by comparing the true TFCE-enhanced *t*-statistic to its empirical null distribution of null TFCE-enhanced *t*-statistics. Finally, we applied the Benjamini-Yekutieli correction to these *p*-values to control the FDR at 0.05.

For correlation analyses between encoding- and retrieval-evoked theta synchrony, we performed a one-sample *t*-test on the within-subject Pearson correlation coefficient values against a null mean of zero. We computed the correlation between synchronous theta encoding and retrieval networks by correlating contact-level hubness scores — i.e., average synchronization to all other contacts — across contacts. Contacts not located in the 80 standard ROIs, such as contacts localized to white matter or the ventricles, were excluded. To take into account the intrinsic neural variability between behavioral events that limits the maximal theoretical correlation between encoding- and retrieval-evoked activity, we sought a benchmark correlation value by conducting a split-half correlation analysis on retrieval events. For each analyzed session, successful–unsuccessful retrieval event pairs were randomly split into two halves, synchrony values and hubness scores were calculated as aforementioned separately for each half, and hubness scores were correlated across contacts.

In the cases of the slow and fast theta comparison and the encoding–retrieval correlation analysis, we have argued in favor of the null hypothesis rather than merely failing to reject the alternative. Because traditional decision-theoretic notions of statistical significance require either rejecting the null hypothesis or declining to make any statistical decision, we turn to the Bayes factor. Bayes factors correspond to a likelihood ratio that can provide evidentiary support for either hypothesis, and the Jeffreys-Zellner-Siow Bayes factor that we employ imposes minimal prior information by incorporating a Cauchy distribution on the standardized effect size and the Jeffreys prior for variance (Rouder et al., 2009).

To analyze the relationship between spectral power and phase synchrony during successful memory in both encoding and retrieval, we quantified the difference in mean power between successful and unsuccessful memory events for each contact with the Cohen’s *d* measure. We used Cohen’s *d* because we were interested in whether the same regions that exhibit large effects of memory-related power changes also exhibit large effects of memory-related synchrony changes. However, between-condition changes in spectral power are less physiologically meaningful if there is greater variability within each condition, and a standardized effect measure not only accounts for the variability of the observations within each condition, but also remains commensurable with the findings of other studies that analyze different sets of behavioral events with different distributional properties. Hence, we measure memory-related power changes with Cohen’s *d* values, a measure widely used in the field to report changes in spectral power and analogous physiological signals. Excluding contacts not localized to one of the 80 standard ROIs, we correlated Cohen’s *d* values with each contact’s hubness score across contacts within subject, and performed a one-sample *t*-test against a null mean of zero on the Pearson correlation coefficients for hypothesis testing.

### Data and code availability

All analyzed datasets, comprising the FR1, catFR1, and pyFR datasets of the Penn Computational Memory Lab, are available online for public use through OpenNeuro (https://memory.psych.upenn.edu/Data). The code used to process and analyze the data and generate the figures is publicly available at https://github.com/adityamrao/RaoEtal24.

## Results

### Synchronous theta networks during memory retrieval

Analyzing data from 382 patients performing a delayed verbal free recall task, we computed inter-trial phase synchrony at 3–8 Hz during the encoding of items (250–1250 ms after word onset) and during the periods immediately preceding spontaneous recall (–1000–0 ms before vocalization). To identify theta synchrony effects specific to stronger memory function, we contrasted subsequently recalled items with not-recalled items presented during the encoding phase, and correctly recalled words with matched deliberation periods (Methods) during the retrieval phase. Replicating previous results in the literature (Solomon et al., 2017; Solomon et al., 2019), we observed increased theta synchrony throughout the whole brain both for successful encoding (*p* = 8.5 × 10^−4^) and for successful retrieval (*p* = 2.3 × 10^−15^). Comparing correct recalls and incorrect recalls, or intrusions, in a subset of 167 subjects for whom at least 10 matched pairs of correct recalls and intrusions were available (see “Methods”) also revealed an effect of theta phase synchrony (*p* = 0.0027). Theta synchrony therefore distinguishes successfully encoded items from more poorly encoded items, memory retrieval from failure to retrieve any memory, and correct recalls from false memories (Figure 2A).

**Figure 2:**
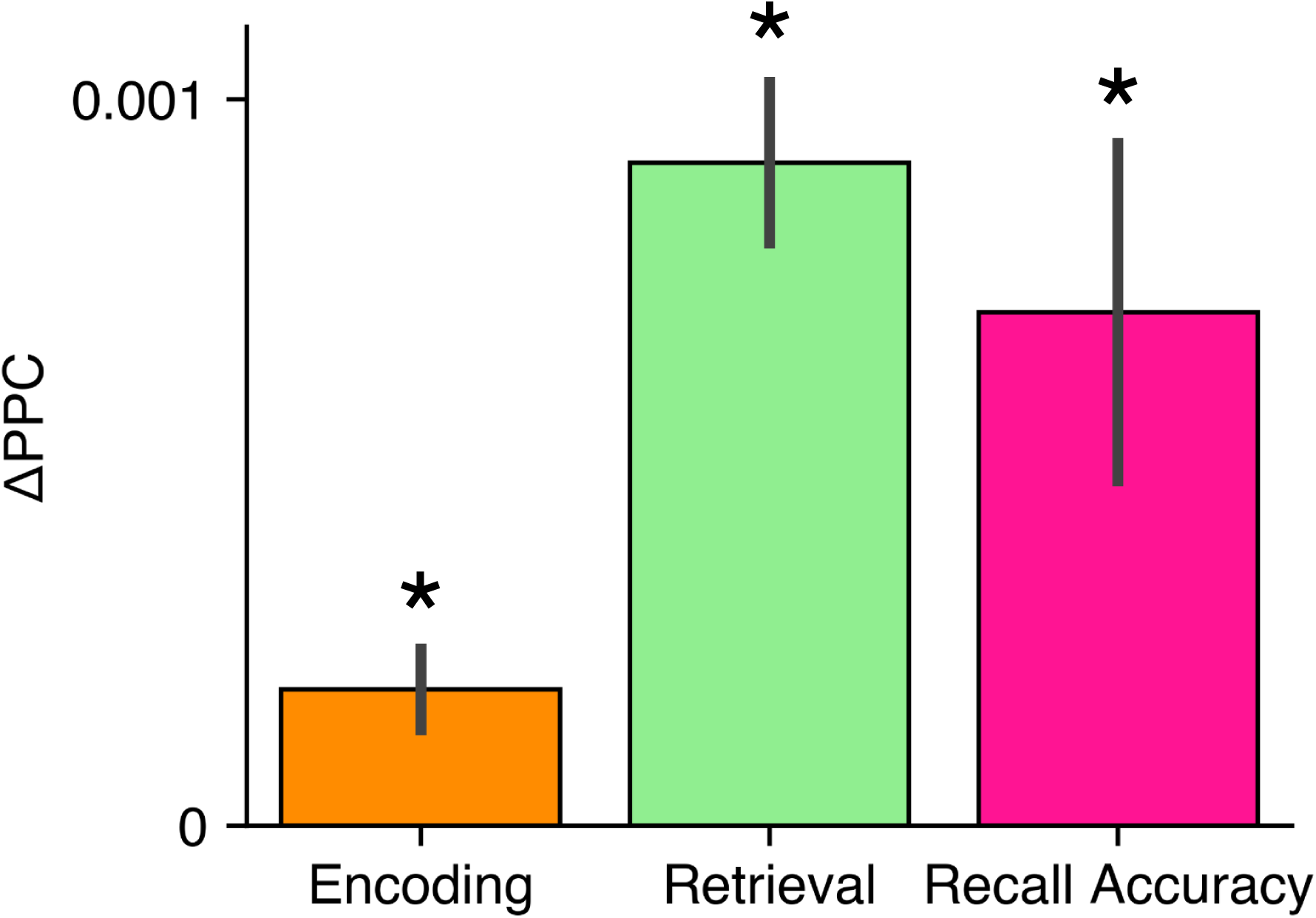
Whole-brain theta synchrony effect, equivalent to the difference in PPC between success-ful memory events and unsuccessful memory events, for encoding, retrieval, and recall accuracy (mean ± SEM across subjects). Asterisk indicates statistical significance at α = 0.05. Figure 2–1 shows the distribution of effects across subjects.

Global theta synchronization during successful memory function might arise from a distributed increase in synchrony among a diffuse set of connections across many brain regions, or from the action of particular regions, or hubs, that strongly synchronize with the rest of the brain. As previous studies have reported network hubs for successful memory encoding, we now sought to determine the hubs of the synchronous theta activity that precedes spontaneous recall. Following the notion of a hub as a node central to a network’s global topology, maintaining widespread connections (Sporns, 2014), we considered a region a hub if its mean synchronization value with all other regions in the brain was reliably positive across subjects. We evaluated 80 ROIs throughout the brain for hubness, controlling the false discovery rate at *Q*_FDR_ = 0.05 in our hypothesis testing. Although we failed to detect any hubs of the encoding network, we detected 33 synchronous hubs and zero desynchronous hubs in the retrieval network, and one synchronous hub and zero desynchronous hubs in the recall accuracy network. Consistent with the theoretical and empirical importance of the hippocampus in human memory, the one synchronous hub found in the recall accuracy network was the left hippocampus. Figure 3 illustrates these hubs and the connections that exhibit the strongest effects (Xia et al., 2013). The retrieval network hubs inhabit diverse regions of the brain, including the medial and lateral temporal lobes, the lateral and medial frontal lobes, and the parietal lobe. Hence, a widespread and distributed synchronous theta network characterizes successful memory retrieval.

**Figure 3:**
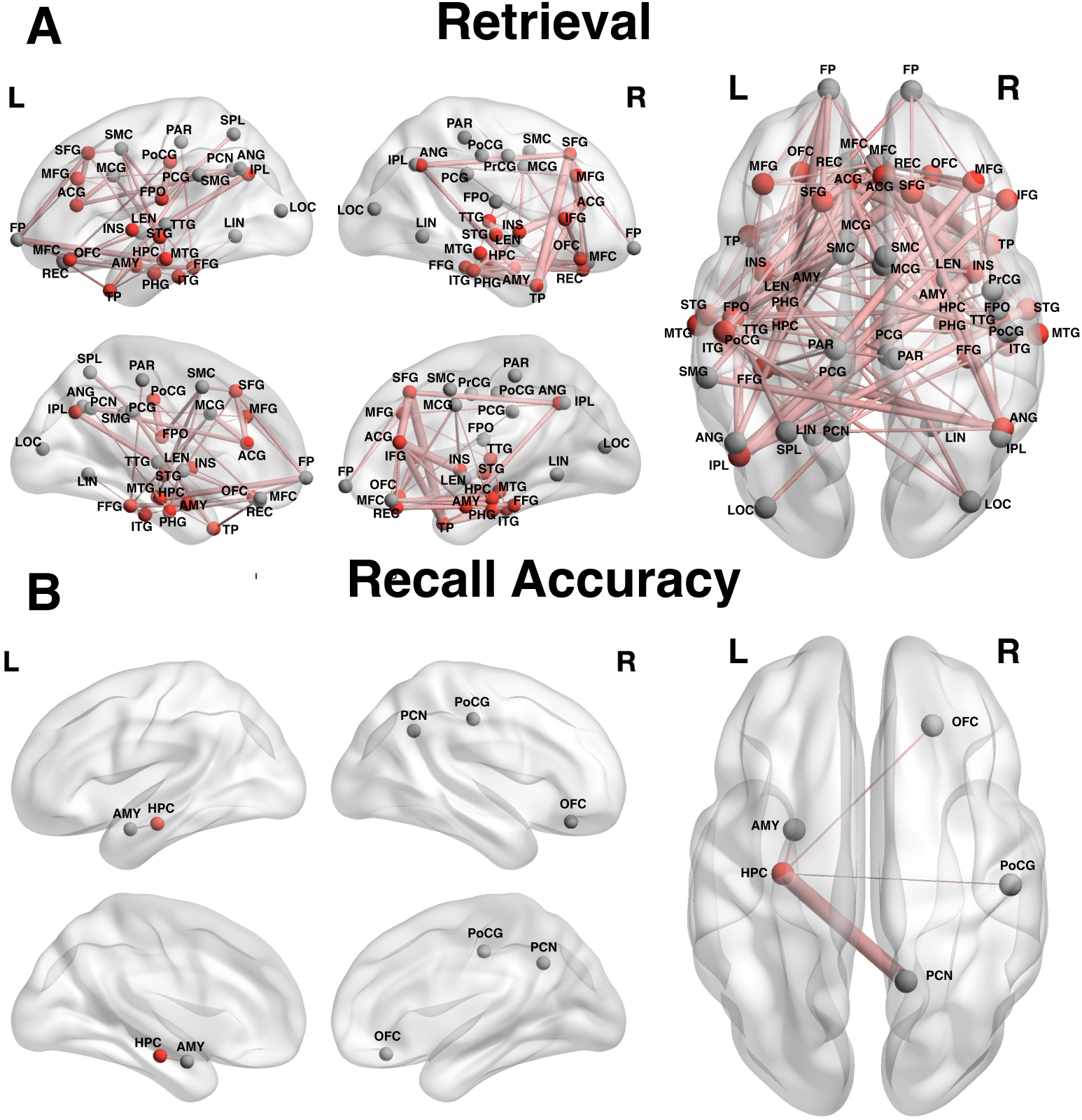
Network hubs and their strongest connections for the retrieval and recall accuracy contrasts. Figure 3–1 enumerates all analyzed region labels and abbreviations. These networks were generated for visualization by plotting the network hubs determined by the hub analysis and for each hub, the five connections with the highest subject-average ΔPPC values, including self-connections and after removing all connections with less than 7 subjects’ worth of data. Positive hubs are colored in red and regions that are not hubs are colored in gray. Width of connections corresponds to strength (|ΔPPC|). Positive connections are colored in red. No negative hubs were identified and hence no negative connections are displayed.

### Role of MTL and hippocampus in synchronous theta networks

Having established that brain-wide increases in theta synchrony mark the imminent recall of previously studied items, we next asked whether the medial temporal lobe (MTL) is a stronger hub than other brain regions implicated in memory: in particular, the lateral temporal cortex (LTC), prefrontal cortex (PFC), and parietal lobe. Although studies of memory have long signaled the special importance of the medial temporal lobe to episodic memory (Staresina et al., 2012; Solomon et al., 2019), prior investigations of the electrophysiology of memory retrieval have identified some of the strongest neural correlates in the lateral temporal and frontal cortices (Burke et al., 2015; Kragel et al., 2017). We hypothesized that the MTL was a stronger hub of the synchronous theta network of memory than the LTC, PFC, or parietal lobe. In comparing the MTL to these lobe-level ROIs, we analyzed the MTL’s mean connectivity to all regions but the subregions of the lobe with which it was compared, and the compared lobe’s mean connectivity to all regions but the subregions of the MTL. The encoding network exhibited no such differential effects (LTC: *t*_299_ = −1.4, *p*_FWE_ = 0.46, PFC: *t*_267_ = 0.83, *p*_FWE_ = 0.80, parietal: *t*_243_ = 0.84, *p*_FWE_ = 0.80), nor did the recall accuracy network (LTC: *t*_132_ = −0.10, *p*_FWE_ = 1.0, PFC: *t*_117_ = 1.0, *p*_FWE_ = 0.91, parietal: *t*_106_ = 0.30, *p*_FWE_ = 1.0). In the retrieval network, the MTL did exhibit stronger global synchrony than the parietal lobe (*t*_243_ = 4.3, *p* = 8.4 × 10^−5^) but we did not find such an effect for the LTC (*t*_299_ = 0.95, *p*_FWE_ = 0.50) or for the PFC (*t*_267_ = 1.1, *p*_FWE_ = 0.50).

Next, we looked within the MTL to the hippocampus, which may maintain interregional connections crucial to memory-related neural activity. Hippocampus–PFC connections are of considerable interest in the functional connectivity literature (Eichenbaum, 2017), and specifically cited for a role in exerting strategic control over memory function, such as selecting the context-appropriate memory during memory search. We examined hippocampal connections both to the dorsolateral PFC (Edin et al., 2009; Anderson et al., 2016; Oehrn et al., 2018), located in the middle frontal gyrus, and to the medial frontal cortex, including the MFC proper and the anterior cingulate gyrus (Euston et al., 2012; Preston and Eichenbaum, 2013; Morici et al., 2022). Controlling the family-wise error rate across these two regions, we did not find an effect of memory-evoked theta synchrony between the hippocampus and dlPFC in the encoding (*t*_194_ = 0.08, *p*_FWE_ = 0.94), retrieval (*t*_194_ = 1.2, *p*_FWE_ = 0.23), or recall accuracy (*t*_81_ = 0.32, *p*_FWE_ = 1.0) networks. Likewise, we did not find evidence for theta synchronization between the hippocampus and mPFC in the encoding (*t*_95_ = 2.0, *p*_FWE_ = 0.11), retrieval (*t*_95_ = 2.1, *p*_FWE_ = 0.08) or recall accuracy (*t*_34_ = 0.64, *p*_FWE_ = 1.0) networks.

### Stronger memory-related increases in slow theta synchrony than fast theta synchrony

The previous analyses describe retrieval-related theta synchrony in diverse brain regions by averaging effects across the canonical 3–8 Hz theta band. However, research on human verbal and spatial memory has associated distinct low and high theta components with different aspects of memory (Lega et al., 2012; Jacobs, 2014; Goyal et al., 2020; Rudoler et al., 2023). In light of the heterogeneity of spectral correlates of memory across the theta band, we asked whether global theta synchrony effects differed between low and high theta frequencies (Figure 4A). Because of the frequency domain leakage arising from Morlet wavelet convolution, we took 3 Hz and 8 Hz to represent low and high theta, respectively. During successful retrieval, regions throughout the brain tended to synchronize both at 3 Hz (*p* = 3.3 × 10^−11^) and at 8 Hz (*p* = 0.0068), but reliably more so at 3 Hz (*p* = 2.5 × 10^−5^). On the other hand, in encoding and recall accuracy, there was a statistically significant effect at 8 Hz (encoding: *p* = 0.0098; recall accuracy: *p* = 0.0096) but not at 3 Hz alone (encoding: *t*_381_ = 1.4, *p* = 0.15; recall accuracy: *t*_166_ = 1.3, *p* = 0.21). Neither the encoding network nor the recall accuracy network exhibited a difference between slow and fast theta (encoding: *t*_381_ = −0.42, *p* = 0.67; recall accuracy: *t*_166_ = −0.41, *p* = 0.68). Indeed, whereas the Bayes factor for the retrieval comparison strongly favors the alternative hypothesis (*B*_01_ = 0.0035), the Bayes factors for the encoding (*B*_01_ = 22.5) and recall accuracy (*B*_01_ = 15.0) comparisons strongly support the null hypothesis.

**Figure 4:**
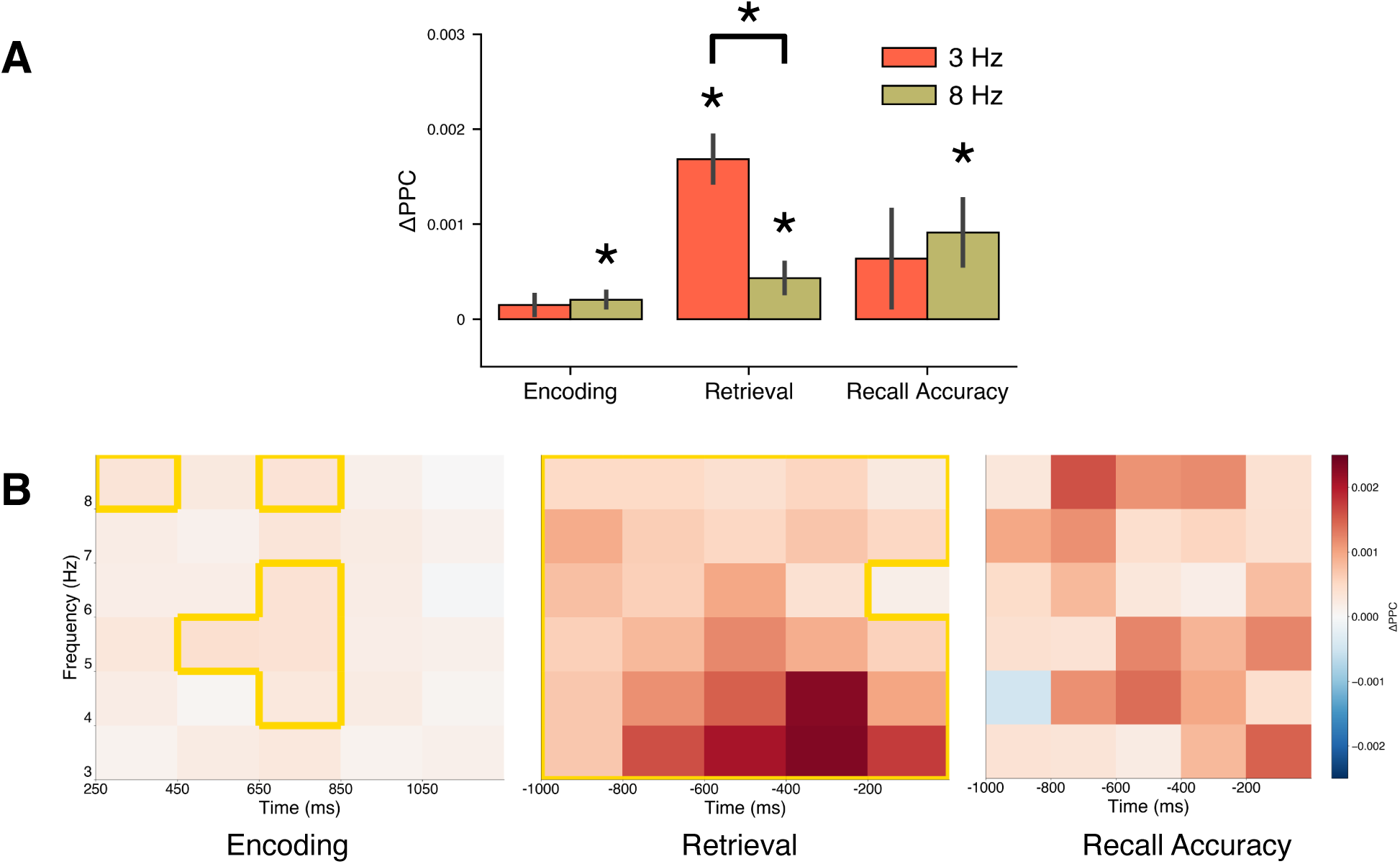
(A) Mean whole-brain connectivity at 3 Hz and 8 Hz for encoding, retrieval, and recall accuracy networks (mean ± SEM across subjects). Asterisk indicates statistical significance at α = 0.05. Figure 4–1 shows the distribution of effects across subjects. (B) Time-frequency plots for encoding, retrieval, and recall accuracy whole-brain theta synchrony effects. Clusters of time-frequency pixels with statistically significant effects are outlined with a yellow border (see “Methods”).

Figure 4B visualizes the time courses of the theta synchrony effects at each theta frequency. Except for a single time-frequency pixel in the recall accuracy contrast, these plots show a consistent association between successful memory and theta synchrony across the relevant time window and the theta band for each behavioral contrast.

### Comparison of encoding and retrieval networks

Having identified the regional distribution of the memory retrieval network, we asked whether it coincided with the synchronous theta network that underlies the encoding of subsequently recalled items. Although prior work has described a common core of cortical regions that are activated during memory encoding and retrieval (Kragel et al., 2017), and shown that spectral patterns during item recall reinstate those present during the same item’s encoding (Manning et al., 2011), studies have also reported differing patterns of theta synchronization during successful encoding and retrieval within the MTL (Solomon et al., 2019). Hence, it is unclear whether widespread theta connectivity of a region during encoding corresponds to similar activation during retrieval.

Correlating hubness between the encoding and retrieval networks across all electrode contacts within subjects revealed a small correlation of *r* = 0.036 ± 0.008 (*p* = 3.0 × 10^−6^). Because neural variability between behavioral events at different times attenuates the maximal observable correlation, we contextualized this finding with a split-half correlation analysis on the retrieval events (“Methods”), obtaining a benchmark correlation value of *r* = 0.053 ±0.006. We considered a similar comparison between the encoding and recall accuracy networks, obtaining a correlation value of *r* = 0.020 ± 0.012 (*p* = 0.089) and a benchmark correlation value of *r* = 0.013 ± 0.009, suggesting a limited ability to detect correlations at the level of contacts between the hubness associated with different sets of events (Figure 5). Bayes factors calculated from the paired-samples *t*-statistic support the hypothesis that the encoding–retrieval and encoding–recall accuracy correlation values do not differ from their benchmarks (encoding–retrieval: *B*_01_ = 4.9, *t*_381_ = −1.8; encoding–recall accuracy: *B*_01_ = 15.0, *t*_166_ = 0.41). Despite the low observed correlation values, perhaps due to the limited quantity of analyzable within-subject recall events, the similarity of the empirical correlation values to the benchmark values established by the split-half correlation analyses suggests that the correlation between the encoding network and the retrieval or recall accuracy network may achieve the noise-imposed upper bound, rather than a lack of relationship between underlying patterns of functional connectivity during encoding and retrieval.

**Figure 5:**
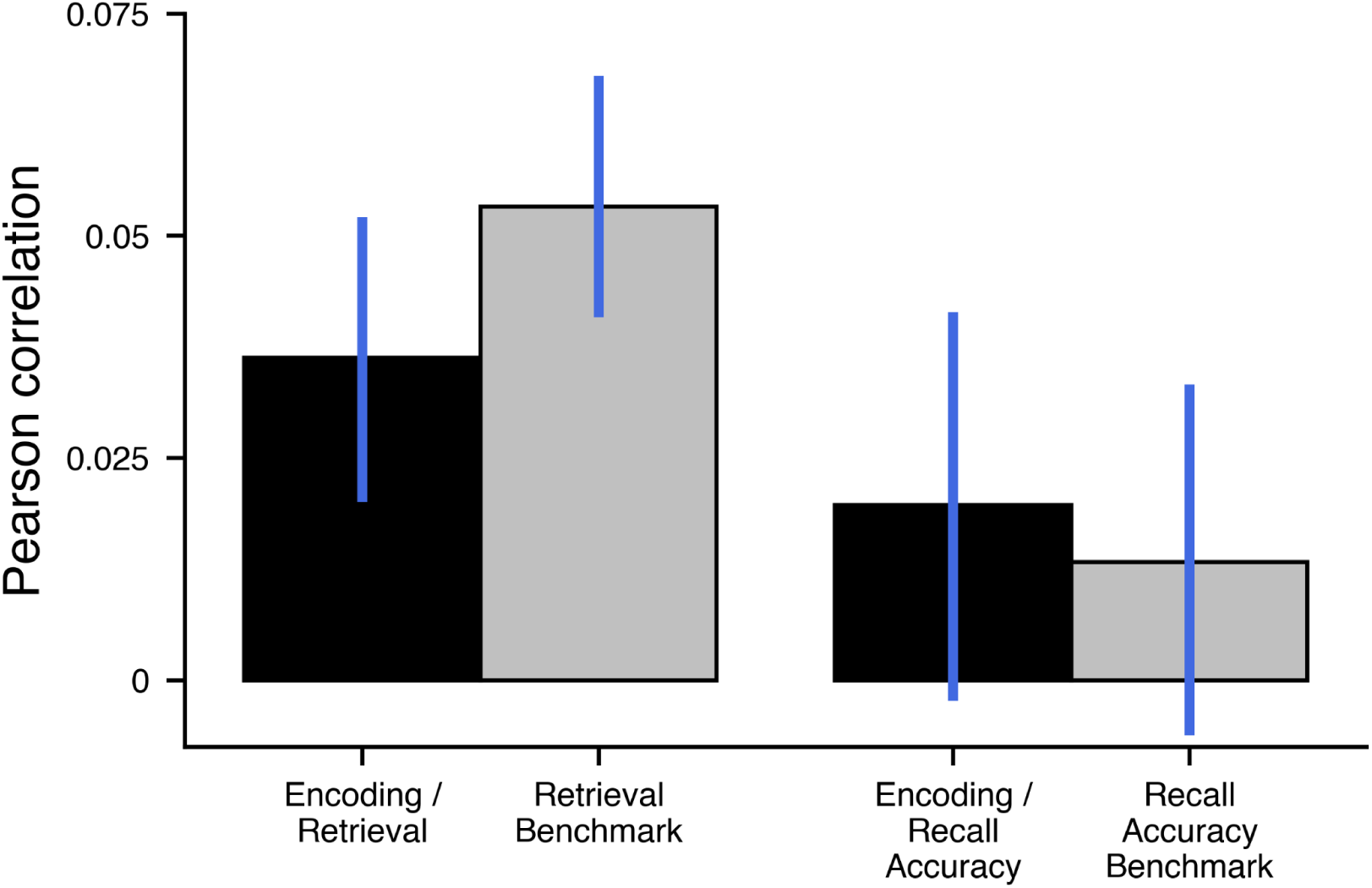
Pearson product-moment correlation (mean ± SEM across subjects) between encoding hubness and retrieval or recall accuracy hubness, and split-half Pearson correlation benchmark for retrieval and recall accuracy hubness. Figure 5–1 shows the distribution of effects across subjects.

### Power–synchrony relationship

Having detailed the whole-brain network of theta synchrony that distinguishes successful retrieval from unsuccessful memory search, we sought to establish its relation to other spectral correlates of memory function. Modulation of theta power also underlies memory retrieval (Burke et al., 2014; Kragel et al., 2017; Rudoler et al., 2023), with broadband low-frequency power decreases and narrowband low theta increases accompanying periods of successful spontaneous recall in comparison to periods of unsuccessful memory search. In addition, theta coherence positively mediates stimulation-evoked increases in theta power (Solomon et al., 2018). We therefore examined how memory-related changes in theta power and theta synchrony covary throughout the brain (Figure 6). Correlating a given region’s change in theta power with its hubness value, or mean brain-wide theta synchrony effect, across contacts for each behavioral contrast revealed a positive correlation between theta synchrony and power effects in the encoding (*r* = 0.26, *p* = 2.5 × 10^−141^), retrieval (*r* = 0.26, *p* = 9.2 × 10^−126^), and recall accuracy (*r* = 0.21, *p* = 1.5 × 10^−45^) behavioral contrasts. Given the earlier findings of stronger memory-evoked synchrony at 3 Hz than at 8 Hz in the retrieval network, we tested for differences in the power–synchrony correlation between 3 Hz and 8 Hz. We found that in recall accuracy, the correlation at 3 Hz, *r* = 0.18, was stronger than the correlation at 8 Hz, *r* = 0.15 (*p* = 0.026), but the differences in the encoding analysis (*p* = 0.63; 3 Hz: *r* = 0.24; 8 Hz: *r* = 0.24) and the retrieval analysis (*p* = 0.19; 3 Hz: *r* = 0.22; 8 Hz: *r* = 0.21) were not statistically significant. In light of findings in the literature of memory-related coupling between theta phase and gamma amplitude (Belluscio et al., 2012; Lega et al., 2015), we also examined the relationship between memory-evoked theta synchrony with memory-evoked gamma power. We found a weak but statistically significant correlation across contacts between gamma power and theta synchrony hubness in the encoding network (*r* = −0.015, *p* = 0.012) and no statistically significant correlation in the retrieval (*p* = 0.51) and recall accuracy (*p* = 0.88) networks.

**Figure 6:**
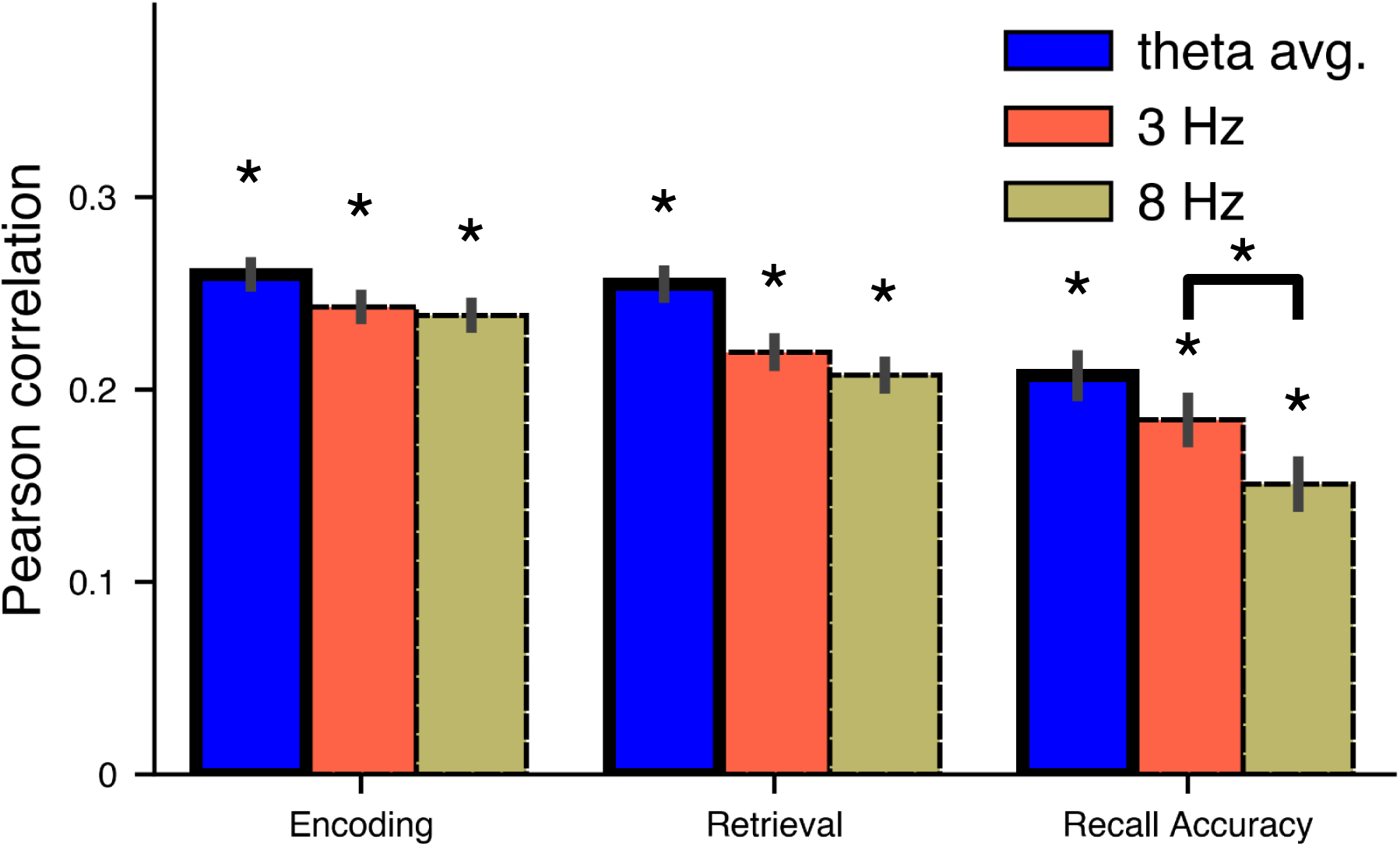
Correlation between spectral power and region-averaged phase synchrony (“hubness”) effects across electrode contacts (mean ± SEM across subjects) of the encoding, retrieval, and recall accuracy networks, with effects analyzed averaged across the theta band, at 3 Hz only, or at 8 Hz only. Asterisk indicates statistical significance at α = 0.05. Figure 6–1 shows the distribution of effects across subjects.

## Discussion

We aimed to characterize the whole-brain synchronous theta networks of successful memory retrieval, leveraging our study’s large dataset comprising recordings from 44,058 intraparenchy-mal and depth electrodes across 382 subjects. We found a marked pattern of whole-brain theta synchrony accompanying successful memory retrieval, and distinguishing the imminent recall of correct items from that of intrusion errors. The synchronous theta networks identified during retrieval spanned widespread brain regions, centering on the prefrontal cortex and lateral and medial temporal lobes, and in agreement with a crucial role for the hippocampus in human memory, we also identified the left hippocampus as a hub of the recall accuracy network. We found reliably greater retrieval-evoked theta synchrony at 3 Hz than at 8 Hz when comparing correct recalls to deliberation periods, but not for the encoding or recall accuracy contrasts. Having demonstrated brain-wide theta synchrony increases during both successful memory encoding and retrieval, we asked whether similar networks underlie both effects. We found reliable but modest correlations between encoding and retrieval networks, but these correlations did not differ from a split-half correlation benchmark. Thus, our analyses are not inconsistent with the hypothesis of encoding– retrieval neural similarity. Finally, we link increases in theta synchrony during successful memory at sites in the brain to higher theta power for both slow and fast theta. While studies of theta phase synchrony in cognition often center on the connectivity of a handful of regions, such as the hippocampus, these results highlight the role of distributed, brain-wide theta synchrony in memory function. They also raise the possibility that phase synchrony modulates the spectral power changes that are of widespread interest in the cognitive electrophysiology literature.

We found that retrieval is marked by brain-wide theta phase synchrony, with network hubs throughout the brain. In the recall accuracy contrast, where our statistical power was more limited, we detected a single hub, the left hippocampus. Solomon et al. (2017) reported a whole-brain theta synchrony effect of retrieval. However, the hubs and connections that constituted the functional networks behind memory retrieval remained unknown, and the patterns of theta synchrony associated with correct recalls in comparison to intrusions had not been investigated. Here, we show that memory retrieval rests on distributed connectivity throughout the brain, and that brain-wide synchrony increases in the left hippocampus characterize the accuracy of the recall. Indeed, prior work analyzing low-frequency spectral power, although more broadly than the theta band, has shown that hippocampal activity predicts recall accuracy (Herz et al., 2023). However, despite the theoretical importance of the MTL and the hippocampus to human memory, we were unable in any of the behavioral contrasts to detect hubness differences between the medial temporal lobe and the prefrontal or lateral temporal cortices, or specific connections between the hippocampus and prefrontal cortex. Further work is necessary to understand the specific contributions of these regions to functional networks of memory in the brain.

We also observed stronger retrieval-related whole-brain synchrony in slow theta than in fast theta, in agreement with a growing recognition of distinct low and high theta bands in human electrophysiology (Lega et al., 2012; Watrous et al., 2013; Jacobs, 2014; Goyal et al., 2020; Rudoler et al., 2023). For example, Watrous et al. (2013) showed that slow and fast theta were differentially associated with spatial recall and temporal recall, respectively, in a forced-choice spatial and temporal retrieval paradigm. Here, we show that in a retrieval–deliberation contrast in episodic free recall, slow theta more strongly drives the synchrony effects associated with better memory function.

After describing retrieval-related synchrony effects, we considered the extent to which the neural activity associated with memory retrieval recapitulates the activity associated with memory encoding. Although phase synchrony is a widely studied neural correlate of cognition, few studies have compared phase synchrony effects across memory encoding and retrieval. However, because similarity of retrieval-related neural activity to encoding-related activity supports the theoretical notion that memory retrieval rests on the reinstatement of the context present during encoding, this line of investigation is important. Hubness across recording sites displayed a relatively low correlation between the encoding and retrieval networks. An earlier study on theta power changes during encoding and retrieval suggested that theta power increases associated with successful encoding and retrieval are uncorrelated across sites in the brain (Lega et al., 2012). Moreover, Kragel et al. (2017), comparing low-frequency spectral power effects between successful encoding and retrieval, found some overlap alongside regional activations unique to each contrast, with much more widespread activation in retrieval. Hence, they proposed that encoding and retrieval processes activate a common core of regions while differing in patterns of interregional connectivity. More recently, a study of hippocampal connectivity observed that theta synchrony patterns specific to memory function appeared to differ between encoding and retrieval (Choi et al., 2020). While our results are not inconsistent with these earlier studies that have compared correct recalls to matched deliberations, our benchmark analysis indicates a low capacity to detect correlations between functional connectivity effects from distinct samples of behavioral events from our limited data.

We also report a positive correlation between a region’s changes in brain-wide theta synchrony and theta power associated with successful encoding and retrieval and accurate recall. Little is known about whether phase coupling mediates changes in power, and if so, how. A previous study related low-frequency coherence to theta power evoked by direct electrical stimulation, alluding to the existence of a relationship between baseline phase synchrony and transient power changes (Solomon et al., 2018). Other research on low-frequency power and phase synchrony correlates of memory formation in a paired-associates learning task found increased 2–4 Hz phase synchrony among regions that exhibited 2–4 Hz power increases during successful memory formation (Haque et al., 2015), but did not systematically investigate the correlation between memory-related changes in global phase synchrony and power across all recorded regions within subjects. Our results significantly extend these suggestions by demonstrating a systematic positive relationship between power and phase synchrony, in both slow and fast theta. Since broadband low-frequency activity decreases during successful encoding and retrieval, one possibility is that regions that show power changes associated with successful memory tend to be distinct from those that exhibit stronger phase synchrony during retrieval. Another is that narrowband power effects form the basis for the power–synchrony correlation. During successful encoding and retrieval, narrowband slow theta power increases, and narrowband fast theta power decreases (Rudoler et al., 2023). In this case, regions that exhibit increases in slow theta phase synchrony also exhibit the memory-related increases in slow theta power. Meanwhile, regions that exhibit fast theta phase synchrony effects of successful memory are less likely to exhibit memory-related narrowband fast theta power decreases. In part, this proposal echoes the hypothesis of Herweg et al. (2020), which aimed to reconcile LFA decreases with theta synchrony increases during successful memory, but that review of the literature on memory-related theta effects did not consider the distinction between slow and fast theta and their opposing narrowband power effects.

Our study offers several avenues for future investigation. First, efforts to unify and replicate disparate results in the literature will enhance our knowledge and theories about functional connectivity. For instance, our inability to detect hubs of the encoding network contrasts with the reporting of dozens of hubs throughout the brain in Solomon et al. (2017). Furthermore, studies might continue to explore experimental paradigms beyond verbal free recall, the experiment for which our intracranial dataset is large enough to make strong inferences. For example, recognition memory may differ in its neural correlates from free recall (Merkow et al., 2014). Recognition memory studies have shown that participants may recognize words that they do not spontaneously recall, and conversely may also recall words that they do not recognize as seen (Kahana et al., 2005; Ozubko et al., 2021). Therefore, recognition memory represents a related and important experimental setting to which to extend analyses of neural activity and functional connectivity. Finally, our study’s methodology, which aggregates sparse and inconsistent electrode montages across subjects, and presents different random selections of word items to different participants, complicates deeper theoretical modeling and analysis of how functional connectivity varies across and within subjects under different parameters. The sources of intersubject and intrasubject variability in functional connectivity and in the neural correlates of cognition more generally thus remain a key target for subsequent research.

These findings indicate that diffuse, global theta synchrony, particularly in slow theta, characterizes successful memory, including encoding, retrieval, and recall accuracy. Understanding the functional connectivity correlates of human memory may improve accounts of the interactive roles of brain regions in memory, lead to subtler descriptions of diseases that impair memory, and even serve as a guide to targeted therapeutic interventions for memory — for example, with electrical stimulation (Ezzyat et al., 2023). In future work, a richer picture of the functional networks of human memory will benefit from relating functional connectivity results across experimental behavioral paradigms and brain recording modalities and from characterizing sources of variability in neural activity associated with cognition and behavior.

## References

Adamovich-Zeitlin R, Wanda PA, Solomon E, Phan T, Lega B, Jobst BC, Gross RE, Ding K, Diaz-Arrastia R, Kahana MJ (2021) Biomarkers of memory variability in traumatic brain injury. Brain Communications 3:fcaa202 Number: 1.

Addante RJ, Watrous AJ, Yonelinas AP, Ekstrom AD, Ranganath C (2011) Prestimulus theta activity predicts correct source memory retrieval. Proceedings of the National Academy of Sciences, USA 108:10702–10707.

Anderson MC, Bunce JG, Barbas H (2016) Prefrontal–hippocampal pathways underlying inhibitory control over memory. Neurobiology of Learning and Memory 134:145–161.

Avants BB, Epstein CL, Grossman M, Gee JC (2008) Symmetric diffeomorphic image registration with cross-correlation: evaluating automated labeling of elderly and neurodegenerative brain. Medical Image Analysis 12:26–41.

Belluscio MA, Mizuseki K, Schmidt R, Kempter R, Buzsáki G (2012) Cross-frequency phase– phase coupling between theta and gamma oscillations in the hippocampus. Journal of Neuroscience 32:423–435.

Benchenane K, Peyrache A, Khamassi M, Tierney PL, Gioanni Y, Battaglia FP, Wiener SI (2010) Coherent theta oscillations and reorganization of spike timing in the hippocampal-prefrontal network upon learning. Neuron 66:921–936.

Benjamini Y, Krieger AM, Yekutieli D (2006) Adaptive linear step-up procedures that control the false discovery rate. Biometrika 93:491–507 Number: 3.

Burke JF, Merkow M, Jacobs J, Kahana MJ, Zaghloul K (2015) Brain computer interface to enhance episodic memory in human participants. Frontiers in Human Neuroscience 8:1055.

Burke JF, Sharan AD, Sperling MR, Ramayya AG, Evans JJ, Healey MK, Beck EN, Davis KA, Lucas TH, Kahana MJ (2014) Theta and high–frequency activity mark spontaneous recall of episodic memories. Journal of Neuroscience 34:11355–11365.

Buzsáki G (2002) Theta oscillations in the hippocampus. Neuron 33:325–340.

Choi K, Bagen L, Robinson L, Umbach G, Rugg M, Lega B (2020) Longitudinal Differences in Human Hippocampal Connectivity During Episodic Memory Processing. Cerebral Cortex Communications 1:tgaa010 Number: 1.

Clouter A, Shapiro KL, Hanslmayr S (2017) Theta Phase Synchronization Is the Glue that Binds Human Associative Memory. Current biology: CB 27:3143–3148.e6 Number: 20.

Desikan R, Segonne B, Fischl B, Quinn B, Dickerson B, Blacker D, Buckner RL, Dale A, Maguire A, Hyman B, Albert M, Killiany N (2006) An automated labeling system for subdividing the human cerebral cortex on MRI scans into gyral based regions of interest. NeuroImage 31:968–80.

Dykstra AR, Chan AM, Quinn BT, Zepeda R, Keller CJ, Cormier J, Madsen JR, Eskandar EN, Cash SS (2012) Individualized localization and cortical surface-based registration of intracranial electrodes. NeuroImage 59:3563–3570.

Edin F, Klingberg T, Johansson P, McNab F, Tegnér J, Compte A (2009) Mechanism for top-down control of working memory capacity. Proceedings of the National Academy of Sciences 106:6802–6807 Number: 16.

Eichenbaum H (2000) A cortical-hippocampal system for declarative memory. Nature Reviews. Neuroscience 1:41–50.

Eichenbaum H (2017) Prefrontal–hippocampal interactions in episodic memory. Nature Reviews Neuroscience 18:547–558 Number: 9.

Euston D, Gruber A, McNaughton B (2012) The Role of Medial Prefrontal Cortex in Memory and Decision Making. Neuron 76:1057–1070 Number: 6.

Ezzyat Y, Kragel JE, Solomon EA, Lega BC, Aronson JP, Jobst BC, Gross RE, Sperling MR, Worrell GA, Sheth SA, Wanda PA, Rizzuto DS, Kahana MJ (2023) Functional and anatomical connectivity predict brain stimulation’s mnemonic effects. Cerebral Cortex p. bhad427.

Fell J, Klaver P, Elfadil H, Schaller C, Elger CE, Fernandez G (2003) Rhinal-hippocampal theta coherence during declarative memory formation: interaction with gamma synchronization? European Journal of Neuroscience 17:1082–1088.

Fell J, Ludowig E, Rosburg T, Axmacher N, Elger C (2008) Phase-locking within human mediotemporal lobe predicts memory formation. NeuroImage 43:410–419.

Fell J, Axmacher N (2011) The role of phase synchronization in memory processes. Nature Reviews Neuroscience 12:105–118 Number: 2.

Fischl B, van der Kouwe A, Destrieux C, Halgren E, Ségonne F, Salat DH, Busa E, Seidman LJ, Goldstein J, Kennedy D, et al. (2004) Automatically parcellating the human cerebral cortex. Cerebral Cortex 14:11–22.

Foster BL, Kaveh A, Dastjerdi M, Miller KJ, Parvizi J (2013) Human retrosplenial cortex displays transient theta phase locking with medial temporal cortex prior to activation during autobiographical memory retrieval. The Journal of Neuroscience 33:10439–10446.

Goyal A, Miller J, Qasim SE, Watrous AJ, Zhang H, Joel M. Stein I CS, Gross RE, Willie JT, Lega B, Lin JJ, Sharan A, Wu C, Sperling MR, Sheth SA, McKhann GM, Smith EH, Catherine S, Jacobs J (2020) Functionally distinct high and low theta oscillations in the human hippocampus. Nature Communications 11.

Haque RU, Wittig JH, Damera SR, Inati SK, Zaghloul KA (2015) Cortical low-frequency power and progressive phase synchrony precede successful memory encoding. The Journal of Neuroscience 35:13577–13586.

Herz N, Bukala BR, Kragel JE, Kahana MJ (2023) Hippocampal activity predicts contextual misattribution of false memories. Proceedings of the National Academy of Sciences 120:e2305292120 Number: 40.

Holm S (1979) A simple sequentially rejective multiple test procedure. Scandinavian journal of statistics pp. 65–70.

Howard MW, Kahana MJ (1999) Contextual variability and serial position effects in free recall. Journal of Experimental Psychology: Learning, Memory, and Cognition 25:923–941.

Jacobs J (2014) Hippocampal theta oscillations are slower in humans than in rodents: implications for models of spatial navigation and memory. Philosophical Transactions of the Royal Society B: Biological Sciences 369:20130304.

Kahana MJ (2012) Foundations of Human Memory Oxford University Press, New York, NY.

Kahana MJ, Rizzuto DS, Schneider A (2005) Theoretical correlations and measured correlations: Relating recognition and recall in four distributed memory models. Journal of Experimental Psychology: Learning, Memory, and Cognition 31:933–953.

Kahana MJ (2020) Computational models of memory search. Annual Review of Psychology 71:107–138.

Kragel JE, Ezzyat Y, Sperling MR, Gorniak R, Worrell GA, Berry BM, Inman C, Lin JJ, Davis KA, Das SR, Stein JM, Jobst BC, Zaghloul KA, Sheth SA, Rizzuto DS, Kahana MJ (2017) Similar patterns of neural activity predict memory function during encoding and retrieval. NeuroImage 155:60–71.

Lachaux JP, Rodriguez E, Martinerie J, Varela FJ (1999) Measuring phase synchrony in brain signals. Human Brain Mapping 8:194–208.

Lega B, Burke JF, Jacobs J, Kahana MJ (2015) Slow theta-to-gamma phase amplitude coupling in human hippocampus supports the formation of new episodic memories. Cerebral Cortex 26:268–278.

Lega B, Jacobs J, Kahana M (2012) Human hippocampal theta oscillations and the formation of episodic memories. Hippocampus 22:748–761.

Manning JR, Polyn SM, Baltuch G, Litt B, Kahana MJ (2011) Oscillatory patterns in temporal lobe reveal context reinstatement during memory search. Proceedings of the National Academy of Sciences, USA 108:12893–12897.

Merkow MB, Burke JF, Stein JM, Kahana MJ (2014) Prestimulus theta in the human hippocampus predicts subsequent recognition but not recall. Hippocampus 24:1562–1569.

Mölle M, Marshall L, Fehm HL, Born J (2002) Eeg theta synchronization conjoined with alpha desynchronization indicate intentional encoding. The European Journal of Neuroscience 15:923–928.

Morici JF, Weisstaub NV, Zold CL (2022) Hippocampal-medial prefrontal cortex network dynamics predict performance during retrieval in a context-guided object memory task. Proceedings of the National Academy of Sciences 119:e2203024119 Number: 20.

Oehrn CR, Fell J, Baumann C, Rosburg T, Ludowig E, Kessler H, Hanslmayr S, Axmacher N (2018) Direct Electrophysiological Evidence for Prefrontal Control of Hippocampal Processing during Voluntary Forgetting. Current Biology 28:3016–3022.e4 Number: 18.

Ozubko JD, Sirianni LA, Ahmad FN, MacLeod CM, Addante RJ (2021) Recallable but not recognizable: The influence of semantic priming in recall paradigms. Cognitive, Affective, & Behavioral Neuroscience 21:119–143.

Preston A, Eichenbaum H (2013) Interplay of Hippocampus and Prefrontal Cortex in Memory. Current Biology 23:R764–R773 Number: 17.

Roberts BM, Clarke A, Addante RJ, Ranganath C (2018) Entrainment enhances theta oscillations and improves episodic memory. Cognitive neuroscience.

Rouder JN, Speckman PL, Sun D, Morey RD, Iverson G (2009) Bayesian t tests for accepting and rejecting the null hypothesis. Psychonomic Bulletin & Review 16:225–237 Number: 2.

Roux F, Parish G, Chelvarajah R, Rollings DT, Sawlani V, Hamer H, Gollwitzer S, Kreiselmeyer G, Ter Wal MJ, Kolibius L, Staresina BP, Wimber M, Self MW, Hanslmayr S (2022) Oscillations support short latency co-firing of neurons during human episodic memory formation. eLife 11:e78109.

Rudoler JH, Herweg NA, Kahana MJ (2023) Hippocampal theta and episodic memory. Journal of Neuroscience 43:613–620.

Rutishauser U, Ross I, Mamelak A, Schuman E (2010) Human memory strength is predicted by theta-frequency phase-locking of single neurons. Nature 464:903–907.

Siapas A, Lubenov E, Wilson M (2005) Prefrontal phase locking to hippocampal theta oscillations. Neuron 46:141–151.

Sigurdsson T, Stark KL, Karayiorgou M, Gogos JA, Gordon JA (2010) Impaired hippocam-pal–prefrontal synchrony in a genetic mouse model of schizophrenia. Nature 464:763–767 Number: 7289.

Smith SM, Nichols TE (2009) Threshold-free cluster enhancement: Addressing problems of smoothing, threshold dependence and localisation in cluster inference. NeuroImage 44:83–98.

Solomon EA, Stein JM, Das S, Gorniak R, Sperling MR, Worrell G, Inman CS, Tan RJ, Jobst BC, Rizzuto DS, Kahana MJ (2019) Dynamic theta networks in the human medial temporal lobe support episodic encoding and retrieval. Current Biology 29:1100–1111.

Solomon EA, Kragel JE, Gross RE, Lega BC, Sperling MRW G, Sheth SA, Zaghloul KA, Jobst BC, Stein JM, Das SR, Gorniak R, Inman CS, Seger S, Rizzuto DS, Kahana MJ (2018) Medial temporal lobe functional connectivity predicts stimulation-induced theta power. Nature Communications 9:4437.

Solomon EA, Kragel JE, Sperling MR, Sharan AD, Worrell GA, Kucewicz MT, Inman CS, Lega BC, Davis KA, Stein JM, Jobst BC, Zaghloul KA, Sheth SA, Rizzuto DS, Kahana MJ (2017) Widespread theta synchrony and high-frequency desynchronization underlies enhanced cognition. Nature Communications 8:1704.

Solomon EA, Lega BC, Sperling MR, Kahana MJ (2019) Hippocampal theta codes for distances in semantic and temporal spaces. Proceedings of the National Academy of Sciences 116:24343–24352 Number: 48.

Sporns O (2014) Contributions and challenges for network models in cognitive neuroscience. Nature Neuroscience 17:652–660.

Squire L, Zola-Morgan S (1991) The medial temporal lobe memory system. Science 253:1380–1386.

Staresina BP, Henson RN, Kriegeskorte N, Alink A (2012) Episodic reinstatement in the medial temporal lobe. Journal of Neuroscience 32:18150–18156.

Vinck M, Lima B, Womelsdorf T, Oostenveld R, Singer W, Neuenschwander S, Fries P (2010) Gamma-phase shifting in awake monkey visual cortex. Journal of Neuroscience 30:1250.

Watrous AJ, Tandon N, Conner CR, Pieters T, Ekstrom AD (2013) Frequency-specific network connectivity increases underlie accurate spatiotemporal memory retrieval. Nature Neuroscience 16:349–356.

Wixted J, Squire L (2011) The medial temporal lobe and the attributes of memory. Trends in Cognitive Sciences 15:210–217.

Xia M, Wang J, He Y (2013) BrainNet viewer: a network visualization tool for human brain connectomics. PLoS One 8:e68910.

Yushkevich PA, Pluta JB, Wang H, Xie L, Ding SL, Gertje EC, Mancuso L, Kliot D, Das SR, Wolk DA (2015) Automated volumetry and regional thickness analysis of hippocampal subfields and me-dial temporal cortical structures in mild cognitive impairment. Human Brain Mapping 36:258–287.

