## Extended Data for "Synchronous theta networks characterize successful memory retrieval"

Figure 2–1

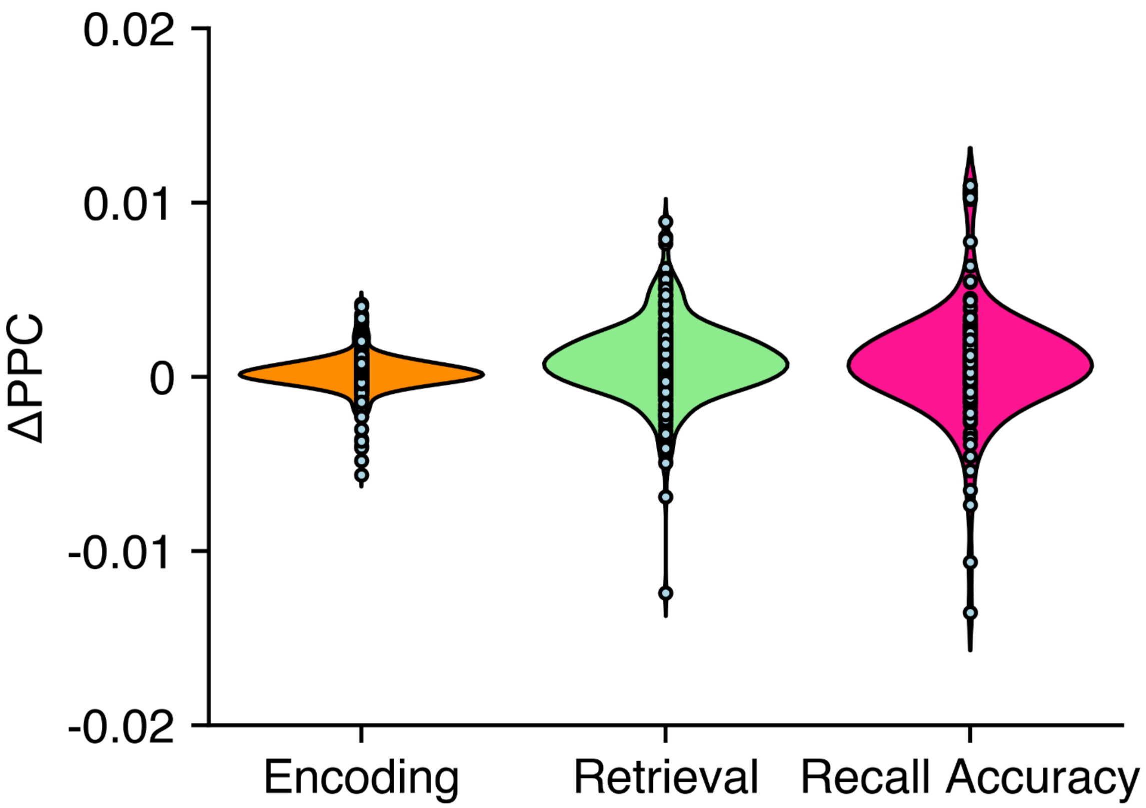


Whole-brain theta synchrony effect for encoding, retrieval, and recall accuracy. Width of the violin indicates density of the distribution, and individual subject data points are plotted along the violin.

Figure 3–1

| **Label** | **Region** | **Label** | **Region** |
|---|---|---|---|
| AMY | Amygdala | MFG | Middle frontal gyrus |
| ANG | Angular gyrus | MTG | Middle temporal gyrus |
| ACG | Anterior cingulate gyrus | OFC | Orbitofrontal cortex |
| BF | Basal forebrain | PAR | Paracentral lobule |
| CAL | Calcarine cortex | PHG | Parahippocampal gyrus |
| CAU | Caudate nucleus | PoCG | Postcentral gyrus |
| CUN | Cuneus | PCG | Posterior cingulate gyrus |
| FP | Frontal pole | PrCG | Precentral gyrus |
| FPO | Frontoparietal operculum | PCN | Precuneus |
| FFG | Fusiform gyrus | REC | Rectal gyrus |
| HPC | Hippocampus | SCA | Subcallosal area |
| IFG | Inferior frontal gyrus | SFG | Superior frontal gyrus |
| IPL | Inferior parietal lobule | SPL | Superior parietal lobule |
| ITG | Inferior temporal gyrus | STG | Superior temporal gyrus |
| INS | Insula | SMC | Supplementary motor cortex |
| LOC | Lateral occipital cortex | SMG | Supramarginal gyrus |
| LEN | Lentiform nucleus | TP | Temporal pole |
| LIN | Lingual gyrus | THA | Thalamus |
| MFC | Medial frontal cortex | TTG | Transverse temporal gyrus |
| MCG | Middle cingulate gyrus | VD | Ventral diencephalon |

Figure 4–1

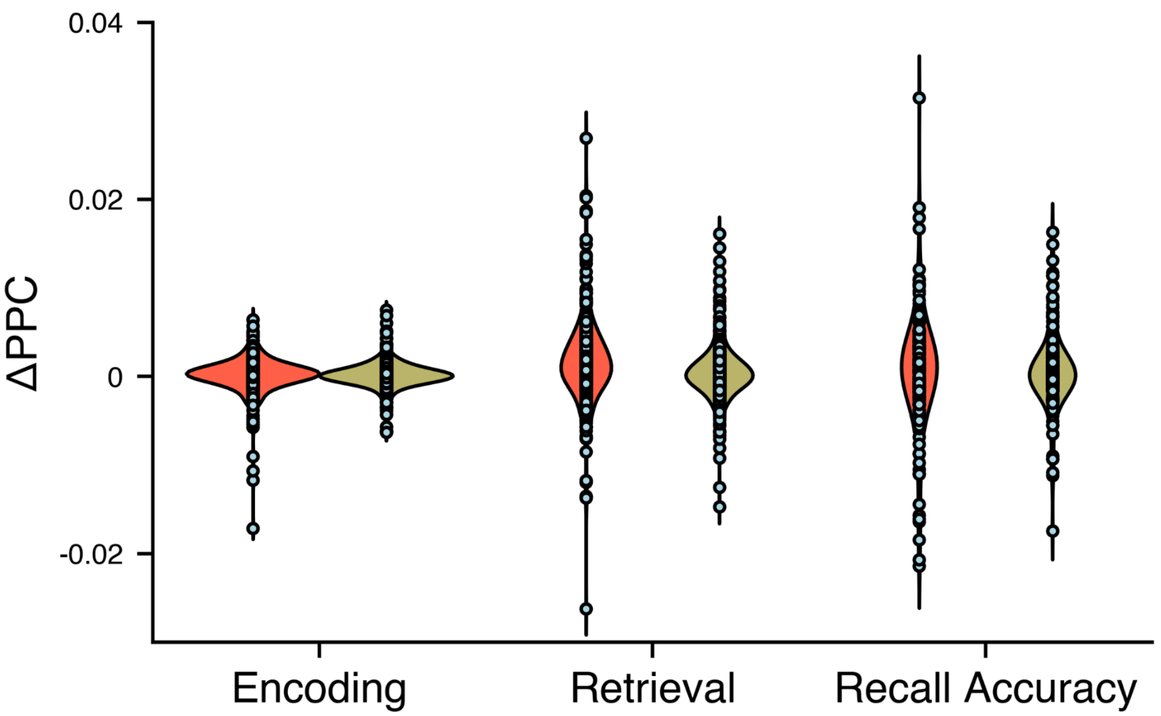


Mean whole-brain connectivity at 3 Hz and 8 Hz for encoding, retrieval, and recall accuracy networks. Width of the violin indicates density of the distribution, and individual subject data points are plotted along the violin.

Figure 5–1

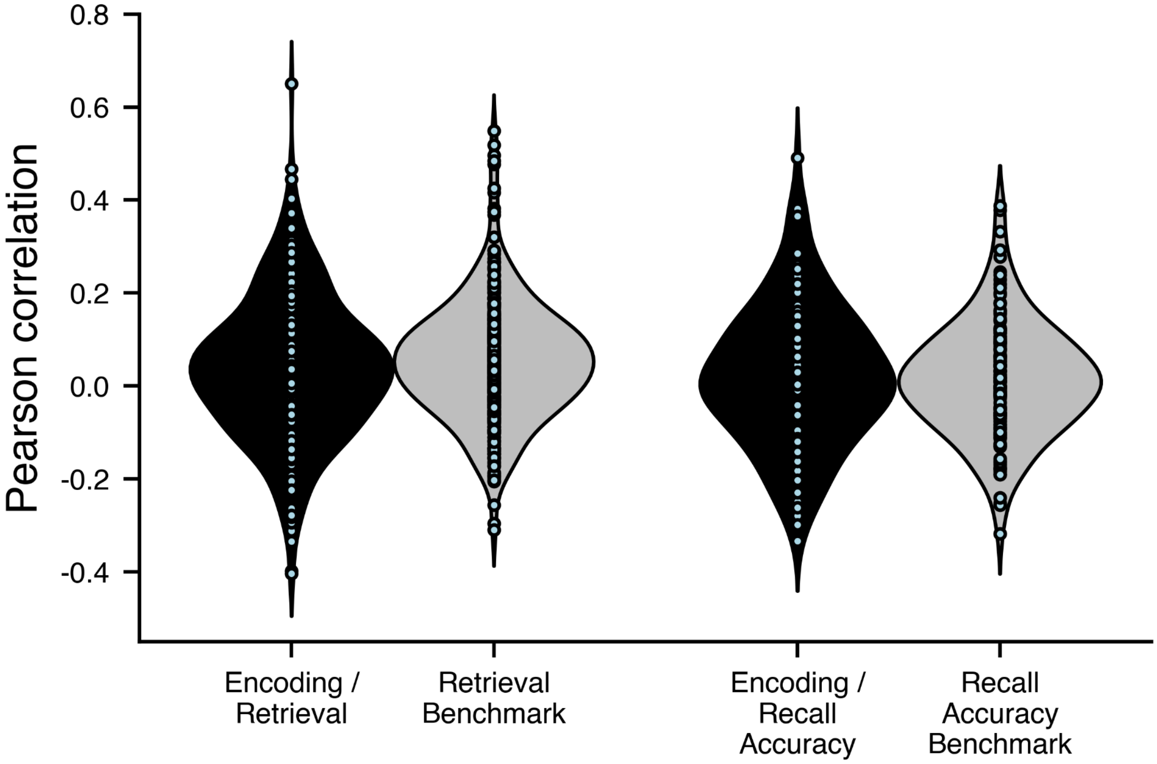


Pearson product-moment correlation between encoding hubness and retrieval or recall accuracy hubness, and split-half Pearson correlation benchmark for retrieval and recall accuracy hubness.. Width of the violin indicates density of the distribution, and individual subject data points are plotted along the violin.

Figure 6–1

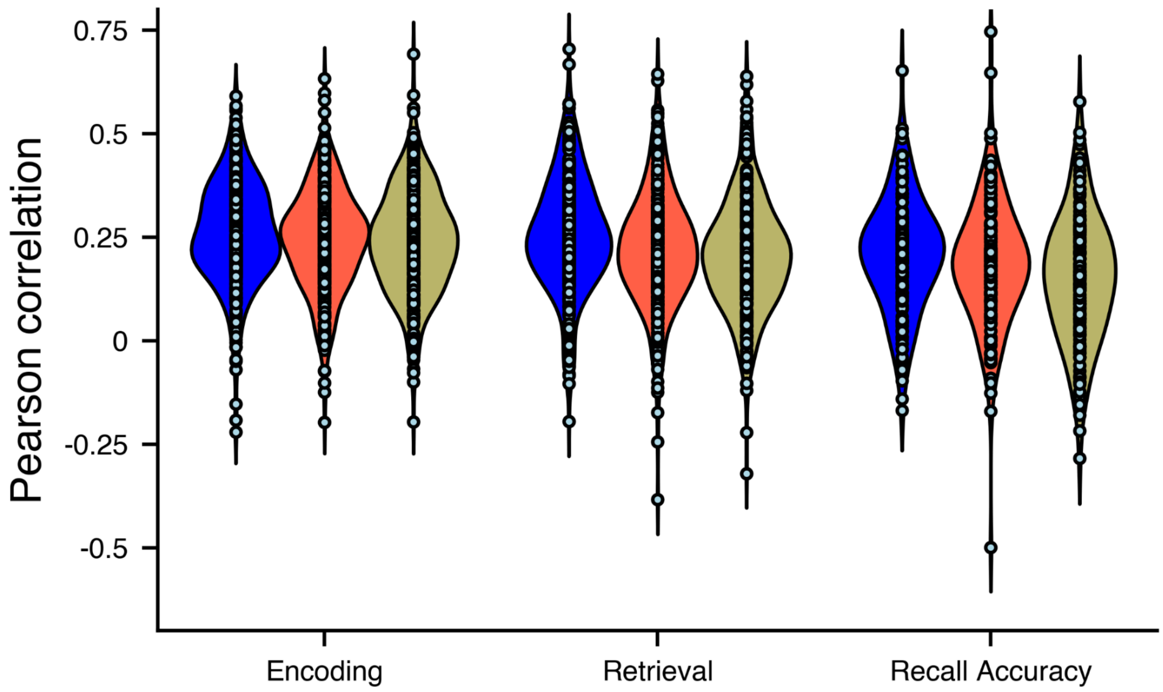


Correlation between spectral power and region-averaged phase synchrony (“hubness”) effects across electrode contacts of the encoding, retrieval, and recall accuracy networks, with effects analyzed averaged across the theta band, at 3 Hz only, or at 8 Hz only. Width of the violin indicates density of the distribution, and individual subject data points are plotted along the violin.
